# From concept to clinic: mathematically informed immunotherapy

**DOI:** 10.1101/027979

**Authors:** Rachel Walker, Heiko Enderling

## Introduction

Mathematical modeling has become a valuable tool in the continued effort to understand, predict and ultimately treat a wide range of cancers in recent years [1,2]. By describing biological phenomena in the concise and formal language of mathematics, it is possible to elucidate key components of complex systems and ultimately develop tools capable of quantifying and predicting system behavior under given conditions. When these tools are applied as a complement to the detailed understanding of cancer biology provided by biological scientists and clinicians, new insights can be gained into the mechanisms and first-order principles of cancer development and control [3].

To date, although mathematical tools have been applied extensively in understanding tumor growth and dynamic interactions between cancer and host, studies involving the theoretical modeling of patient response to treatment and the contribution of such findings to the development of clinically-actionable therapeutic protocols remain strikingly limited. In particular, despite the rising emergence of immunotherapy as a promising cancer treatment, knowledge gained from mathematical modeling of tumor-immune interactions often still eludes application to the clinic. The currently underutilized potential of such techniques to forecast response to treatment, aid the development of immunotherapeutic regimes and ultimately streamline the transition from innovative concept to clinical practice is hence the focus of this review.

## Mathematical oncology

Due to the simplifications inherent in the development of a mathematical model of a complex biological system, early attempts at combining mathematics with biology were often met with skepticism from experimentalists and clinicians. The misconception that simplifications render mathematical models meaningless is still found in some more conservative areas of medical research. Thankfully, the great potential for theoretical models to help unravel highly complex and multifaceted processes without the need for time-consuming experimentation is becoming more widely appreciated [4]. Insights can be gained into mechanisms or relationships that may not be obvious or intuitive, allowing the generation of novel hypotheses on which to base future experimentation.

Formalized descriptions of dynamic processes that have achieved a balance of both simplicity and applicability have contributed greatly to biological research to date. For example, in the field of oncology, tumor growth has been modeled extensively using methods ranging from simple one-equation models featuring logistic, Gompertz or exponential growth functions to detailed 3-dimensional spatio-temporal systems [5-15]. Detailed reviews of tumor growth modeling over the last few decades are plentiful [16-18]. More detailed considerations of invasive morphology have been provided, emphasizing non-linear growth patterns such as fingering and fragmentation [19-24], and invasion models have been developed accounting for phenotypic competition for space and resources in a microenvironment featuring selective forces such as hypoxia and low nutrient concentration [20,21]. Computational models of angiogenesis [25-31], necrosis [19], metastatic processes [21] and stem cell driven growth dynamics and heterogeneity [32-36] have also made valuable contributions to our understanding of cancer biology.

Of equal (or arguably greater) importance, computational modeling can also contribute to the advancement of treatment protocols. Making the transition from initial concept to the implementation of a new treatment protocol in the clinic can be an expensive, laborious and time-consuming process. Phase I trial development traditionally involves simple dose-escalation strategies to identify a maximally tolerated dose based on the observation of toxicity to patients, followed by repeated assessment of adverse side effects in subsequent trials at varying schedules [37]. The great time and cost burden of repeat trials also leads to little time being allocated to updating or improving the protocol in later years after initial demonstration of clinical response or failure. The application of computational tools to provide guidance based on quantitative science may overcome some of these limitations and help to modernize this trial development process, potentially lowering both the physical cost to enrolled patients and the time taken for a protocol to reach the clinic [7].

The primary goal in developing a clinically useful model is to create a tool that can, when provided with a set of initial conditions, reliably reproduce empirical results. When the ability to reproduce clinical or experimental outcomes has been verified, it can then be used to make predictions about previously untested hypothetical scenarios. If said computational tools can model human pathology and physiology it is possible to simulate a range of parameters to represent variations in individual-specific characteristics that correspond to unique “virtual patients” [38,39]. In doing so, a much larger heterogeneous cohort than would be feasible in the clinic can be simulated to predict a myriad of possible outcomes and help forecast potential treatment responses. If these models are carefully parameterized and validated using experimental and clinical data, this could provide a platform for analysis of treatment effectiveness and allow a quick and cost-effective extension of trial populations with limited risk to human patients.

Through such virtual trials, mathematical modeling also has great potential to contribute to the optimization of drug administration schedules. There may often be a wide range of possibilities for dosage, sequencing or duration of treatment which cannot all realistically be tested in the clinic. From a preliminary understanding of the mechanisms involved in patient response, mathematical models can be developed to test any imaginable sequencing option, adjust days or times of treatment, or alter dose distribution within clinically accepted ranges across the duration of the trial. These applications of mathematics remain inadequately utilized when it comes to developing clinical protocols, in particular in the emerging areas of immuno- and combination therapy.

Experimentally and clinically obtained patient-specific parameters, for example individual tumor growth rate or immune response efficacy, can be also incorporated into validated models to optimize outcomes for the particular patient and hence improve overall prognoses. With this capability, modified regimes may also be suggested during treatment to account for individual response, thoroughly personalizing treatment. In addition, the ability to observe tumor response continually is a great benefit of computational simulations of therapy, and can offer supplemental insights into potential tumor volume reduction when radiological imaging is limited to anticipated clinically meaningful time points.

The potential of mathematical modeling to contribute to the field of oncology by improving our understanding of complex biological processes and aiding therapeutic design is vast. Using such methods, disease progression can be more comprehensively understood by bridging the gap between observable clinical outcomes and simulations of the underlying unknown, or unobservable, biological processes [40]. In other areas of medicine such models have already established themselves as valuable tools in paving the way for more efficient clinical trial design. The innovative work of Lasota and Mackey relating to therapy-induced anemia encouraged early models of treatment response [41]. Valuable progress has also been made through the utilization of mathematics in areas such as HIV treatment [42-46]. Despite this, in terms of immunotherapy for malignant cancers, many of the prospective applications in terms of aiding the transition of theoretical knowledge into clinical practice remain a long way from achieving their full potential.

## Immunotherapy

Traditional cancer therapies such as radiotherapy and chemotherapy have had a great deal of clinical success when it comes to decreasing tumor burden and prolonging survival. Despite this, inherent risks such as radiation-induced secondary cancers, healthy tissue toxicity and side effects such as hair loss, weakened immune systems and extreme fatigue are still a concern for many patients and their healthcare providers. As such, a great deal of emphasis has been placed on the development of immunotherapeutic strategies in recent years, which have the potential to limit the negative side effects seen as a result of other therapies when used as an alternative or adjuvant treatment by taking advantage of the body’s own defense mechanisms. For a discussion of the clinical development of the field of immunotherapy and recent clinical trials, the interested reader is directed to [47].

The host immune response to a growing tumor is a highly complex process. One of the critical mechanisms in effective antitumor immunity requires tumor-specific antigens to be recognized by dendritic cells and macrophages which can present this antigen to cytotoxic T lymphocytes capable of mounting an attack on associated tumor cells. However, tumors also take advantage of multiple mechanisms for evading recognition and destruction by the host immune system. For example, by controlling surface molecule expression, tumor cells can dramatically reduce their lysis by cytotoxic T lymphocytes, allowing their continued growth and invasion even in the presence of an adequate immune cell population. This active evasion of attack and elimination by immune cells has been classified as a new and crucial “hallmark of cancer” [48]. As this immune evasion permits tumor growth and metastasis, immune responses to tumor presence are a key target for research and therapeutic advancement.

The goal of immunotherapy is to stimulate the immune system in order to encourage antitumor activity: to boost or restore system ability to fight cancer. Therapeutic modulation of the natural immune response is primarily achieved by enhancing activated effector T cell or immune memory cell activity, augmenting the antigen-presenting capabilities of dendritic cells and chemically heightening tumor immunogenicity and reducing immune-suppression. With the help of immunotherapy, the ultimate goal is to reduce tumor mass and armor the immune system to “remember” the tumor antigens to prevent future recurrence; modern techniques also aim to develop anti-tumor vaccines capable of destroying tumor cells even prior to clinical detection.

Immunotherapy can be a challenging field in which to make clinical progress. Immune responses are often difficult to sustain, and the dynamic and volatile nature of tumor-immune population dynamics can lead to counter-intuitive growth and regression behavior, for example a burst of growth may in fact precede a rapid decline in tumor volume. Small variations in immune response efficacy may also be sufficient to trigger either complete tumor extinction or an explosive period of growth. Additionally, the time taken for cytotoxic effector cell recruitment to initiate significant tumor volume reduction can be greater than the response time of alternative therapies in which regression begins earlier in treatment. This combination of factors can lead to difficulties initiating and maintaining large-scale clinical trials, which may be considered too unpredictable to implement in a clinical setting or may have been prematurely abandoned. Critical hurdles to the advancement of cancer immunotherapy remain, including the complexity of immune escape processes, the currently limited consideration of individual response patterns to cytotoxic agents and immunotherapies and, crucially, the scarcity of collaborative teams of translational researchers dedicated to the field [49]. Past and present challenges facing immunotherapeutic clinical trials have been discussed in detail elsewhere [50-53]. These difficulties can prove to be a great hindrance to the development of novel treatment strategies or their sequencing to harness and exploit the great therapeutic power of the immune system.

To overcome several of these challenges in immunotherapy, quantitative tools can be particularly useful. Computational models, which are able to isolate key mechanisms and simulate spatio-temporal dynamics in the relationship between host immune system and tumor, can provide valuable insights into processes that may be too complex to analyze experimentally. These dynamic interactions can be observed and analyzed extensively through virtual trials *in silico* [38]. Thorough evaluation of potentially counter-intuitive population dynamics and identification of stable or tumor-free equilibrium states under the application of treatment can dramatically enhance and streamline trial development and drug testing procedures.

## Modeling tumor-immune interactions

The self-regulatory nature of the immune system, as well as tumor-immune interactions and the influence of immunotherapy, feature complex and nonlinear spatial and temporal dynamics with significant cellular heterogeneity and extensive signaling and decisionmaking networks [54]. The complexity of cytokine modulation and the large variation in patient-specific immune capabilities cause further complications; ultimately, accounting for every variable involved in tumor-immune interactions is a daunting task. Techniques for mathematical representation of parts of these complex systems to identify overarching principles will be described, along with methods used to incorporate a variety of immunotherapeutic and combination treatments.

In particular, the high level of complexity makes tumor-immune interactions highly amenable to computational modeling by means of differential equations. Such models have been widely applied and tested in the natural sciences for many years [55], and their solution can reach high levels of accuracy when correctly parameterized. Typically, in modeling tumor-immune system interactions and both adaptive and innate immune responses, the dynamics of each population of interest such as lymphocytes or tumor cells can be modeled by a differential equation representing the manner in which said population changes over time. These may feature simple functions for differentiation, growth or proliferation, death, and transport or migration in or out of the compartment. This approach mirrors that found in basic predator/prey approaches to temporal population dynamics in the field of ecology [56].

Simple models of this form featuring the “hunted” tumor cells and the “hunter” immune cell populations and demonstrating extinction, uncontrolled growth and both stable and unstable dormancy can be found in [57-64]. Many other models feature a similar structure, albeit with varying levels of complexity; as many as 15 or 20 coupled differential equations for each cellular subtype or chemical mediator concentration could be utilized to describe immune responses. However, although differential equation models can often be formulated with relative ease, the trade-off between complexity and computational cost leads to a tendency of modelers to utilize more simple, parameterizable models. Insightful summaries of models in tumor-immune interactions can be found elsewhere [65,66].

A general and concise example of the structure of tumor-immune interaction equations can be given by the following pair of ordinary differential equations (ODEs):

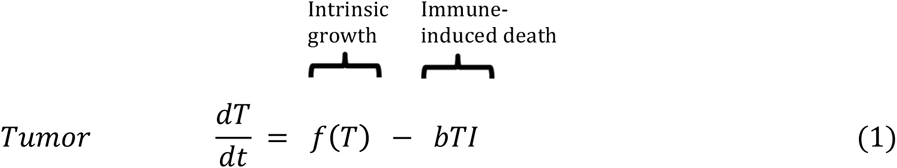

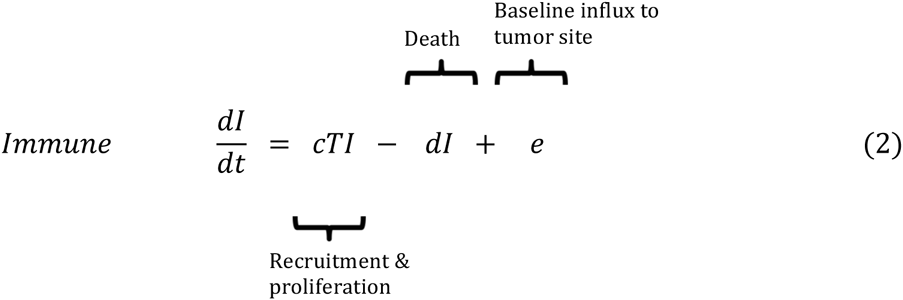

In the example above, populations *T* and *I* represent tumor and immune cell populations, respectively. Function *f(T)* represents net tumor growth; exponential, logistic and Gompertz functions are all commonly used and further detail on these models as well as several alternatives can be found in [67]. A simple schematic of these basic tumor-immune population dynamics is provided in Fig. 1.

**Fig.1.**
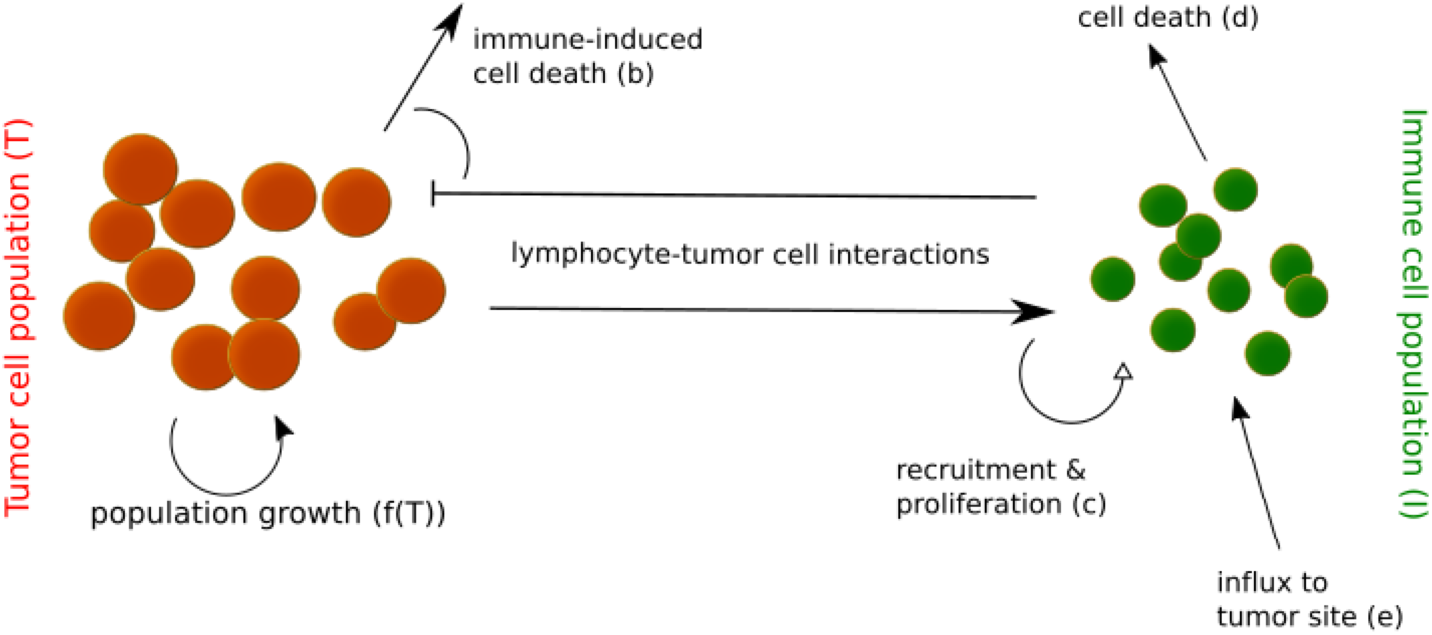
Schematic of tumor-immune dynamics in a basic two-population model. Letters in brackets correspond to parameters in Eqn (1) and (2).

This fundamental structure representing simple lymphocyte-tumor interactions has been extensively adapted to incorporate more complex features of the tumor-immune relationship over the last several decades, including cellular heterogeneity, spatial dynamics, delayed feedback, and cytokine activity and other signaling and modulating factors, to name only a few. Crucially, many of these tumor-immune models could also incorporated immunotherapeutic treatment regimes in order to simulate patient response and predict treatment success. Some of the most insightful of these models of the last few decades will be summarized to facilitate discussion of the contributions and limitations of current research. As the primary emphasis of the present review is on the application of therapy and the associated clinical relevance of model predictions, details of computational solvers will mostly be omitted – for a summary of some of the many numerical methods for approximating solutions to differential equations the unfamiliar reader is directed to, for example, [68]

Cytotoxic effector lymphocytes are commonly modeled as the immune population in the above simple but robust system of ODEs. One of the most widely referenced mathematical descriptions of tumor-immune dynamics was introduced by Kuznetsov et al. and involves modeling the response of CD8+ T cells to an immunogenic tumor and the corresponding kinetics of tumor growth and regression [62]. Above a threshold number of tumor cells growth is able to continue uncontrolled, and below this threshold oscillatory growth and regression is observed, a now widely accepted phenomenon in solid tumor immunoediting [63,64,69-71]. As well as T-lymphocytes, the cytotoxic effects of natural killer (NK) cells on the tumor population can be modeled in a similar manner. NK cells can act as a surveillance effector population; tumors with low antigenicity may experience mainly NK cytotoxic effects whereas those with high antigenicity have been found to primarily experience T cell cytotoxicity [66]. The two independent anti-tumor mechanisms can act with varying intensities, and both cytotoxic effector cell populations may be modeled simultaneously. Accounting for the saturation of effectiveness of CD8+ T cell killing, a strong patient-specific adaptive CD8+ T-lymphocyte response is typically necessary to promote tumor regression and improve the likelihood of treatment success [73].

Despite the key role in the host immune response of cytotoxic effector cells, insights into the behavior of other immune cell populations can also be gained from simple mathematical models. Simulations of interactions between macrophages and T lymphocytes accounting for antigen presentation have demonstrated that the magnitude of the adaptive response can be dramatically increased by the early activation of CD4+ helper T cells, which allows early macrophage accumulation of tumor cell debris and a rapidly mounting immune attack [74,75]. Regulatory T cells can also influence tumor development and dormancy; in tumors featuring low immunogenicity and a high growth rate, effector T cells are able to outcompete regulatory T cells but are still unable to control the tumor [76]. For immunogenic tumors with a slow growth rate, the expansion of effector and regulatory cells is balanced, preventing tumor destruction.

Simple ODE models are useful for considerations of temporal population dynamics, but to thoroughly analyze certain elements of the immune response, spatial variation may need to be taken into account. For example, even in the case of a stable temporal state of dormancy, heterogeneous spatial distributions of both tumor and immune cell populations may be present in immunogenic tumors [64]. Heterogeneous spatial patterning has also been found in the presence of macrophages, which have the potential to affect tumor composition and structural characteristics by altering chemokine activity [77,78]. The effect of such chemical mediators on tumor-immune interactions necessitates a further increased level of model complexity in which reaction or diffusion processes such as the exchange of substances with the local microenvironment may need to be accounted for. For example, nutrient concentration and ECM-degrading enzymes can have substantial effects on tumor growth dynamics [10], and tumor-suppression has been observed when removal of mutant cells by immune cells reaches a critical level dictated by the difference in mitotic rates of the respective cells under the influence of biochemical control mechanisms [79].

Delayed feedback due to transit time or signal delivery can be captured in a simple adaptation of a basic population model. Incorporating the response time of the immune system after recognition of non-self cells using a system of delay differential equations (DDEs) can demonstrate different stability dynamics to a comparable model without delay. Stronger oscillatory behaviors suggestive of the continued regrowth of the tumor can be observed with increasing delay before immune response [58]. Cancer and immune biology also feature stochastic or probabilistic events, presenting a further layer of complexity in tumor-immune modeling. As opposed to whole population dynamics, the action and interaction of discrete and distinguishable individual cells may need to be considered. Each cell follows a distinct set of rules in such models, and individual cell diversity can be accounted for to allow more detailed modeling of cellular-level biological phenomena. Reproduction of experimental observations of Gompertz growth has been achieved in such an agent-based model describing immune surveillance [14].

Considering both chemical species reaction-diffusion processes and cellular level individual-based interactions in a multi-scale hybrid model can allow further insights into patterns of both regression and invasion [80]. A strong immune response with high lymphocyte recruitment and high kill capacity can lead to greater reductions of tumor size [69], and slower nutrient uptake can indicate spatial stability and the development of a substantial necrotic core [19]. In contrast, increased nutrient uptake by tumor cells may demonstrate increased cell motility and morphologic variation in the form of branched growth. Additionally, lower cell adhesion can lead to quicker tumor cell movement to favorable nutrient conditions in which the population can grow [19].

As can be seen even from this brief summary, there exists a broad range of applications of mathematical models to the improved understanding of tumor-immune interactions, a full discussion of which is beyond the scope of the present review. Additional discussion of model types and selection in immunology can be found in [81-83]. With this foundation, the next logical step is to utilize such models to adapt and improve treatment protocols and guide clinical practice.

## Modeling treatment outcomes

Several methods for achieving an immune system boost and improving tumor-specific adaptive immunity have been considered theoretically and in preclinical studies, including the instillation of various cytokines, adoptive cell therapies, vaccine therapies and checkpoint inhibitors. As described previously, making the transition from concept to clinic in the case of immunotherapeutic strategies can be a long and challenging process, and despite extensive research, few such protocols have established themselves as a standard treatment. While monoclonal antibodies and to a lesser extent cytokine-based drugs and checkpoint inhibitors have had some clinical success in treatment of melanoma and lymphomas amongst one or two other specific cancers, the rate of FDA approval of immunotherapeutic treatments lags significantly behind our rapidly advancing knowledge and understanding of the field [84]. This has led to the increased utilization of mathematical models designed to predict the outcomes of hypothetical treatment scenarios to offer insights into the mechanisms underlying the potential success or failure of therapy in terms of tumor remission, dormancy and regrowth, with the intention of guiding future clinical trial design.

The effects of various therapeutic strategies can be modeled by generic cell kill terms or simple adjustment of the fundamental growth, death and interaction parameters of a 2-equation system comparable to that in Eqn. (1) and (2) [61,85]. For example, increasing the immune cell population initial condition to represent the instillation of effector T lymphocytes to a patient, or the imposition of a negative growth function for a tumor cell population to represent conventional chemo-radiation, can demonstrate initial growth acceleration but the ultimate diminishing of the tumor cell population, albeit after clinically unrealistic treatment durations [61]. As clinically verified data to support the many parameters of more complex models are often scarce, such simple complex mathematical models can be robust and insightful, yet for more specific treatments model complexity may be obliged to increase somewhat.

## Adoptive T-cell and cytokine therapies

Tumor-infiltrating lymphocyte (TIL) therapy is a form of adoptive cell transfer (ACT) which involves the clinical extraction of antigen-specific T-lymphocytes from a patient tumor, which can then undergo rapid expansion *ex vivo* by incubation with cytokines and be transfused back into the patient in vast numbers to launch a targeted attack on the tumor cells. Increasing the flow of effector cells into a tumor, or a local increase in the rate of their proliferation, can provide a simple representation of the effects of impulse injections of effector lymphocytes. Although complete tumor destruction may not be possible under such a model, tumor mass can be reduced and tumors may persist in dormant state [86]. Therapies such as ACT have appeared to be more effective in tumor-destruction and side effect limitation than alternative treatments such as T helper cell-stimulating vaccine therapies in recent computational studies [87]. Despite this, only a small reduction in either the probability of a cytotoxic effector cell killing or deactivating a tumor cell, or the rate of influx of these lymphocytes into the tumor vicinity, has been found sufficient to provoke tumor regrowth [86]. It is clear that overcoming lapses in immune response which can lead to the tendency for regrowth is crucial: immune memory boosting in combination with other treatment protocols has demonstrated the potential to help to establish stable dormancy [88].

Immunomodulating cytokines are often utilized both in isolation and in concert with ACT therapies such as TIL to assist with response maintenance. Various cytokines which can mediate both innate and adaptive immunity have undergone extensive clinical and preclinical testing in recent years [89]. Of these, the interleukin family remains one of the most widely studied. IL-2 is primarily responsible for the activation, growth and differentiation of lymphocytes and is one of few cytokine-based treatments to receive FDA approval. When applied in isolation, models have observed the potential for an immune response capable of growing without bounds which could lead to harmful side effects such as capillary leak syndrome in clinical patients [63], although if both tumor antigenicity and the level of IL-2 are high, this side effect may be reduced [90]. However, when both IL-2 and effector lymphocytes are applied together, negative side effects can be minimized while also achieving significant tumor volume reduction or even complete elimination [63,90,91].

IL-21 is additionally capable of directly modulating the volume and function of dendritic, NK and T-lymphocyte cells involved in the immune response. This includes regulating the transition from innate immune responses featuring natural killer cells to adaptive responses featuring activated T-lymphocytes. Enhanced T cell activity without too much inhibition of the NK cell response is required for successful treatment, and this balance may be highly dependent on tumor immunogenicity. In simulations of IL-21 modulation, efficient disease elimination does not occur in immunogenic tumors; only in non-immunogenic tumors can IL-21 stimulate cellular immunity against cancer, and only then if dosage is calculated based on tumor mass at time of administration [92].

Generic immunostimulating therapies such as ACT or cytokine injection can be modeled in less complex models but with the intention of assessing constant, periodic and impulsive instillations [93,94,60] as opposed to stationary treatments. The eradication of tumor cells has been found to depend only on the mean value of the therapy term as opposed to on the shape on the function for realistic therapeutic durations [93], although further evidence of the aforementioned potential for boundless growth of the immune response suggests further tests would be required prior to clinical applicability.

## Dendritic cell vaccines

The augmentation of antigen presenting cell (APC) activity is another way to arm the immune system against tumor cells [95]. Dendritic cells can be loaded with tumor-specific antigens *in vitro,* allowing them to be returned to the patient in a similar manner to that of TIL therapy but now with the goal of presenting the antigen en masse and arming cytotoxic T lymphocytes against tumor cells. In comparing the effects of such dendritic cell vaccines to TIL therapy it has been observed that high numbers of APCs can delay tumor recurrence after regression, whereas high numbers of CTLs can actually have the opposite effect [96]. It is hypothesized that the cytotoxicity of activated lymphocytes to each other may be responsible for this phenomenon, potentially facilitating tumor growth as opposed to hindering it. Alongside the understanding of basic growth and regression dynamics, optimal control theory can be utilized to determine the optimal administration time and dosage of dendritic cells to reduce tumor mass and also minimize the number of injected APCs [97]. Considering both discrete and continuous injections, it can be found that to optimize this form of therapy in the clinic, a high dose initial instillation is required to promptly reduce the tumor, following which smaller doses should continue at approximately even intervals for the duration of treatment.

## Preventative vaccines

In recent years consideration has been given to the possibility of eliminating microscopic initial tumor cell populations prior to clinical detection as an alternative to the treatment of pre-existing, macroscopic tumors. To achieve this would require the development of preventative cancer vaccines which, although still far from clinical applicability and dependent on tumor-specific factors such as cell quantity and mitotic activity, could be a game changing development in the field of immunotherapy. The goal of such therapy is to train or sensitize the immune system, with particular emphasis on cytotoxic T lymphocytes, to identify and destroy tumor cells as soon as possible after initial genesis and prior to their acquisition of immune-suppression or metastatic characteristics. Considering both tumor site and lymph node dynamics under the influence of a vaccine modeled by anti-tumor memory CTLs which proliferate when activated by dendritic cells can suggest the number of cytotoxic T cells that need to be armed against a tumor for an immune memory-based vaccine to be effective [98]. This value may be surprisingly low: a memory lymphocyte population of only 3% or less is found to be sufficient to eradicate a tumor across a wide range of parameters.

## Cancer-specific treatments

Mathematical models of specific cancers and treatments as opposed to more general concepts may incorporate more biologically realistic, detailed characteristics of tumor-immune interactions due to the availability of clinical data with which they can be parameterized, calibrated and validated. Bacillus Calmette-Gue’rin (BCG), a tuberculosis vaccine, is a clinically established treatment for superficial bladder cancer [99]. The standard of care BCG treatment involves one intravesical instillation each week for a period of six weeks, and experiences a clinical response rate of up to 50%-70%. When tumor cells internalize BCG a strong immune response is initiated; the increase in cytokines (such as IL-2) leads to an influx of innate, non-specific effector lymphocytes. When these effector cells reach the site of BCG, they can either degranulate or activate, both of which have the potential to destroy the infected cell and cause bystander tissue trauma, ultimately eliminating the tumor. It is worth noting that an adaptive immune response may also contribute to the success of BCG therapy, given that to achieve the observed clinical response rate the approximate number of bystander cells killed per activated innate cell has been found in simulations to far exceed the natural capacity of the immune system [100].

As this is a therapy already present and effective in clinical practice, computational modeling is a highly useful tool: there exists human patient data from the clinic with which to validate models. Several studies have incorporated BCG therapy into tumor-immune models to gain a deeper understanding of its curative mechanisms and to make suggestions for improvements to standard therapy protocols. For example, it has been found that in pulsed BCG therapy, a decrease in pulse frequency may lead to treatment failure, and an increased dose can generate potentially harmfully large effector cell populations [101,102]. To minimize these risks, a stability condition for the tumor-free state may be defined by upper and lower bounds for the average dose per unit time [102]. BCG is also typically used in an adjuvant capacity post-resection; shortening the time between surgery and the first BCG instillation may be beneficial to patient outcome, as may extending the indwelling time beyond the current standard of care [103]. Furthermore, despite initial computational evidence to the contrary [101,102], patient response may also improve significantly under the combination of both BCG and IL-2 infusion [104].

Malignant gliomas are another prominent area of modeling-based research due to the limited success of existing therapeutic strategies. The administration of membrane glycoprotein T11 target structure (T11TS) as a therapeutic agent has been shown to reverse the immune suppressed state of brain tumor by boosting the functional status of immune cells including macrophages and CD8+ T lymphocytes in animals. Modeling this treatment computationally can predict that treatment with T11TS can allow effector cells of the immune system to overcome blood brain barrier (BBB) impermeability and lead to enhanced phagocytic activity and diminishing of glioma cells [105]. CTL therapy has also been attempted as a treatment of glioma; previous failure of such therapies may be attributable to the administered dose being up to 20x lower than would be required for tumor eradication in this protected region, according to recent simulations of patient response [106]. Additionally, a constant infusion of T lymphocytes may need to be maintained after tumor volume reaches a near stable-state [107].

## Combination therapies

Immunotherapy applied in concert with other therapeutic regimes has led to significant successes in both clinical and pre-clinical trials in recent years [108,109]. Several such regimes have been found to significantly increase the probability of total remission. Despite currently limited utilization for this purpose, mathematical modeling has also proven beneficial in investigating synergy between therapies and optimization techniques for both adjuvant and neo-adjuvant immune-boosting treatments in the few cases of its application to date.

As chemotherapeutic agents are designed to target dividing cells for systemic treatment of advanced cancers, in many patients extreme side effects can occur due to the toxicity of the chemicals used to other, non-cancerous rapidly proliferating cells such as those of the blood, mouth, digestive system, hair and nails. As such, these treatments have more recently been extended to include additional therapies that can reduce the need for such high dosages of toxic agents. In patients with a naturally weak immune response, neither IL-2 immunotherapy nor chemotherapy may be sufficient to trigger substantial tumor regression [110]. However, particularly in patients with a strong immune response, chemotherapy followed by pulsed immunotherapy may stunt growth substantially. Both treatments concurrently was also found to be favorable due to the lower toxicity and greater immune stability, and such combination treatments have been suggested as a promising alternative to monotherapy in other computational studies [111]. A combination of 7 pulsed doses of chemotherapeutic drug along with the injection of 8 x 10^8 activated CD8+ T lymphocytes has been found to lead to a rapid decline in tumor population, with regrowth additionally avoided by the addition of IL-2 treatment. Similar success was achieved under combinations of T cell boosts, IL-2 and cancer vaccines [112,113]. A selection of other combination immune- and chemotherapy treatments have been modeled computationally [114,115]. The main observation from such models to date has been that the synergistic effects of combination treatments are sufficiently beneficial to justify further investigations and implementation of such methods in the clinic on a more regular basis.

During radiation therapy, highly energized particles cause damage to cellular DNA, leading to cell cycle arrest, senescence, and cell death [116]. An ongoing need to reduce damage to normal, healthy tissue and improve clinical outcomes has led to a succession of both clinical and computational attempts to improve and optimize treatment schedules for such therapies both alone and as part of combination treatments. Synergistic combinations of radiation and immunotherapy have shown promise in the clinical setting [108,117-120], yet mathematical models of such regimes are scarce. Despite this, in what may prove to be an important step towards the more effective contribution of mathematics to clinical oncology, a recent study has utilized a model incorporating systemic T cell trafficking to support the hypothesis that local immune activation in an isolated metastatic site followed by targeted radiation can lead to abscopal effects in other metastatic tumors distant to the initially targeted area [121]. Such mathematical modeling may help to identify promising treatment targets on a per-patient basis.

Other models of immunotherapeutic treatments are interspersed throughout the literature. In prostate cancer vaccination therapies, the dosage and interval required to generate stable prostate-specific antigen levels has been found to vary greatly between individuals [122], yet increases in dosage and administration frequency may be able to stabilize disease progression in most patients [123]. IL-12 treatment has the potential to overcome the imbalance between Th1 and Th2 type lymphocytes in melanoma which can lead to a transition from immunosurveillance to immunotolerance, boosting antitumor immune pathways while minimizing harmful side effects [124]. Eleven consecutive doses of siRNA are sufficient to control oscillatory tumor behavior and limit the inhibitory effects on the immune system of TGF-β [125]. Tumor eradication by means of oncolytic virus therapies may require widespread spatial distribution of the virus throughout the tumor, particularly when vascularization has occurred [126]. The list goes on; many treatment protocols are amenable to mathematical modeling. The selection highlighted here is not exhaustive yet emphasizes the important insights into the mechanisms underlying treatment success and failure that can be gained from such computational tools.

## Discussion

Mathematical modeling has made many important contributions to the understanding of tumor growth, dormancy, regression and recurrence in recent years. Many of these models also account for tumor-immune interactions and both the innate and adaptive immune response, and have generally been accepted as valuable tools in aiding the biological and physiological understanding of cancer development. More recently, *in silico* studies have considered the effect of potential immunotherapeutic and combination treatments. Despite providing valuable insights into fundamental mechanisms underlying treatment success and failure, many of the findings of these studies are focused on basic system dynamics and remain far removed from clinical applicability. The observations made in such studies are yet to translate into the clinic and make significant contributions to the development of successful treatment protocols. Also, several significant immunotherapeutic strategies that have undergone preclinical and clinical testing are yet to be considered mathematically, and existing virtual trials may lag behind the most recent innovations in the biological realm due to the lack of sufficient sharable clinical data. The full potential of mathematical modeling to contribute to the improvement of cancer therapy is far from being reached.

The question is, how can this be resolved? Why do examples of the contribution of computational modeling to successful clinical therapy design remain so sparse, and what steps need to be taken to align the capabilities of mathematicians, physicists and computer scientists with the needs of clinicians and patients, to streamline the transition from theoretical concept to clinical practice and improve patient-specific, adaptive immuno- and combination therapies?

A crucial first step in bridging this gap is a greater level of collaboration between those with data, and those with models. The limited availability of valuable, relevant biological data with which to both parameterize and validate mathematical models can be greatly inhibiting. At present, best efforts are made to parameterize numerical models as thoroughly as possible, yet still great difficulty arises in finding recent, relevant and accurate data sources from which to derive them. The continued development of collaborative relationships is critical for understanding underlying concepts, accuracy in assumption-making and the extraction of usable parameters that reflect experimental observations. Several authors have made efforts to validate quantitative models in a human clinical setting (for example [127,128]), but the majority rely on small experimental cohorts of non-human responders, if experimental data is available at all. Conducting data fitting with very limited data points is an additional pitfall. To improve specificity and accuracy in modeling for use in clinical practice much more temporally and spatially resolved clinical data are desperately needed. With improved cross-disciplinary collaborations, patient data could be more effectively collected and utilized to build more efficient model. Biologically informed models can thereafter be tested extensively and authenticated by comparison to actual patient response. In addition, such collaborations can encourage a new era of multi-disciplinary science; theoretical knowledge and experimental techniques should not be utilized in isolation but rather as complementary, synergistic components of successful precision medicine team science [129].

An additional requirement for the improved contribution of mathematical sciences to therapy design is the development of more models able to consider a wide range of doses and delivery schedules for particular immunotherapeutic and combination treatments. To overcome current shortfalls in trial design and to optimize treatment protocols for the quickest improvements in clinical response, a paradigm shift from the use of mathematics for primarily proof-of-concept demonstrations of underlying principles to a combination of both theoretical models and treatment optimization needs to occur. Simulation of many hypothetical regimens and identification of the optimal time and discrete dosing of therapeutic agents to generate the greatest patient response can allow the suggestion of improved protocols for both immunotherapy and combination treatments. Although some examples of more tangible, concrete suggestions for treatment protocols do exist (for example, [130] offers suggestions for improving current standard of care featuring optimization of dose and scheduling to maximize survival, and similar suggestions are made in [106,107,112,124]) such examples still remain sparse in the routine setting [131]. From early considerations of the effect of treatment regimes on patient response [132,133] through to the extensive considerations of optimization of a range of treatments for an equally wide variety of cancers in recent years, the work of Agur, Kronik, Kogan and colleagues continues to pave the way for the advancement of models designed to forecast patient outcomes [134,135]. Others have also begun to consider the simulation of individual treatment responses over large virtual patient cohorts to guide therapy and aid clinicians and innovative models of synergistic therapies can aid the development of novel hypotheses that may have wide reaching impacts in the oncological community such as the optimization of the abscopal effect [121]. However, the value of such prominent works needs to be widely appreciated to encourage further development of clinically motivated biomathematical models that can become commonly utilized by medical practitioners in the advent of immunotherapy.

Based on model predictions of optimal schedule and dose as well as patient-specific disease characteristics, personalized treatment protocols are the highest objective of immunotherapy design to which mathematical modeling has great potential to contribute. To account for the co-evolution of a particular host immune system and tumor, patient-specific inherent immunity must be taken into account. To achieve this goal, clinical data from individual patients needs to be utilized to evaluate patient-specific parameters such as those relating to the natural immune response. Incorporating such parameters in validated models allows the simulation of *personalized* responses to many different hypothetical doses and delivery schedules to identify the optimal regime. This must be a future development of clinical study design: personalized treatments which additionally permit adaptive treatment modification of schedule or dosing to maximize effectiveness for a particular patient and improve clinical outcomes.

## Summary

To allow innovative, progressive treatments to make the transition from concept to clinic, experimentally calibrated and clinically motivated mathematical models are essential. To aid the development of well-informed clinical trials, collaboration between mathematicians, computational modelers, biologists and clinicians is required to allow qualitative hypotheses to take on a quantitative dimension. Such inter-disciplinary science can allow both personalization and optimization of dose and scheduling of immunotherapeutic protocols, both as an independent therapy and in conjunction with other more traditional therapies, to streamline the transition from innovative concept to clinical practice, and improve clinical outcomes for individual patients.

## Acknowledgements

This publication is supported by the Personalized Medicine Award 09-33000-15-03 from the DeBartolo Family Personalized Medicine Institute Pilot Research Awards in Personalized Medicine (PRAPM). We thank Kimberly Luddy, Sotiris Prokopiou and Jan Poleszczuk for fruitful discussions and critical reading of the manuscript.

